# Mitochondrial D-loop DNA analyses of Norway Lobster (*Nephrops norvegicus*) reveals genetic isolation between Atlantic and Mediterranean populations

**DOI:** 10.1101/258392

**Authors:** Jeanne Gallagher, John A. Finarelli, Jónas P. Jonasson, Jens Carlsson

**Affiliations:** School of Biology and Environmental Science/Earth Institute, University College Dublin, Belfield, Dublin 4, Ireland; Marine and Freshwater Research Institute, Skúlagata 4, 101 Reykjavík, Iceland

**Keywords:** Genetic structure, mitochondrial DNA, control region, phylogenetics, Atlantic, Mediterranean, glacial refugia

## Abstract

*Nephrops norvegicus* is a commercially valuable demersal fisheries species. Relatively little is understood about this species’ population dynamics across its distribution with previous mitochondrial and microsatellite studies failing to identify significant population-level differentiation. In this study, sequence variation in the mitochondrial (mtDNA) D-loop was analysed from samples across the distribution range. Analysis of a 375bp fragment of the D-loop revealed significant genetic differentiation between samples from the northeast Atlantic and the East Mediterranean (F_ST_ = 0.107, P<0.001). Tau (τ), theta (θ_0_ and θ_1_) and Fu’s F_s_ values suggest the species spread between 10,500 to 19,000 ybp and subsequently expanded rapidly across the Atlantic.

## Introduction

*Nephrops norvegicus* (Linnaeus, 1758) is a benthic-dwelling, decapod crustacean, which inhabits burrows in patches of soft muddy sediment between approximately 4 to 800m depth (Holthius, 1991; Johnson et al., 2013). The species’ distribution ranges from Iceland and northern Norway, in the North Atlantic Ocean, to Morocco and the Mediterranean in the south (Figure 1) (Maltagliati et al., 1998; Bell et al., 2006; Johnson et al., 2013). *Nephrops norvegicus* is dioecious, with mating occurring following a brief courtship shortly after females moult (Powell and Eriksson, 2013). Females produce between 900-6000 eggs in a brood (Powell and Eriksson, 2013), with dispersal occurring in the larval phase, which can last up to fifty days (Hill, 1990; Dickey-Collas et al., 2000). Survival of the larvae depends on a combination of factors including suitable temperature, food availability and access to suitable substrate (Dickey-Collas et al., 2000; Aguzzi and Sardà, 2008; Pochelon et al., 2009). Upon settling juveniles occupy or create burrows to avoid predation (Powell and Eriksson, 2013). Adult *N. norvegicus* do not migrate or leave their mud-patch at any point (Aguzzi and Sardà, 2008).

**Figure 1.**
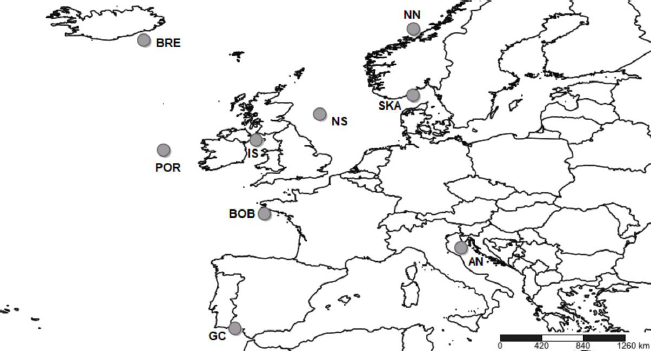
Nine sample locations (n=30 per locality) for *N. norvegicus* mtDNA D-loop analyses. AN: Anacona, BOB: Bay of Biscay, BRE: Breiðamerkurdjúp, GC: Gulf of Cadiz, IS: Irish Sea (West), NN: Northern Norway, NS: North Sea, POR: Porcupine, SKA: Skagerrak

Commonly sold as Norway lobster, Dublin Bay prawn, scampi or langoustine, *N. norvegicus* is a commercially important fisheries species (Thorpe et al., 2000) with the most recent global landing estimates at 54,000 tonnes in 2014 (FAO; Thorpe et al., 2000). Within the EU, 2017 *N. norvegicus* landings are estimated to be worth over €165 million (Marine Institute, 2017). For management purposes the species is currently divided into approximately forty geographic groups, known as functional units (FU’s) and geographical survey areas (GSA’s), across its distribution (Relini et al., 1999; Ungfors et al., 2013).

Effective management relies on accurate and reliable information on how species are distributed over time and space. Current assessment of *N. norvegicus* is largely based on underwater video surveys (Johnson et al., 2013; Marine Institute, 2016). Although the species has a relatively long larval stage (fifty days), the low mobility of adults may increase the vulnerability of stocks to local overfishing relative to other highly mobile commercial species. Commercial fishing has been suggested as the principal driver of population dynamics for the species (Thorpe et al., 2000). Despite the substantial economic value of *N. norvegicus* fisheries, there is limited knowledge of the species’ genetic population structure and whether it aligns with existing functional, biological or management units (Stamatis et al., 2006).

Population genetics has proven highly suited for identifying biological populations by quantifying the connectivity (gene flow/isolation) among them. Population genetics can also assess vital demographic parameters, such as effective population size, evolutionary history and recent demographic expansion (Beissinger and McCullough, 2002). Mitochondrial DNA (mtDNA) has several advantages in population genetic studies. For example, as a maternally-inherited, haploid marker there is a lack of genetic recombination which is ideal for studying deep historical population dynamics (Held et al., 2016). Zane et al. (2000) used single strand conformation polymorphism analyses of mtDNA in populations of Northern krill, *Meganyctiphanes norvegica* (M. Sars, 1857) to reveal at least three distinct populations in the North-East Atlantic. Yuhara et al. (2014) utilised mtDNA Cytochrome c oxidase subunit I (COI) analyses to clarify the genetic diversity and connectivity among local coastal populations of the saltmarsh sesarmid crab, *Clistocoeloma sinense* (Shen, 1933) around the Japanese coastline.

With respect to *N. norvegicus*, allozyme analyses on 110 individuals from one Scottish and two Mediterranean localities (Aegean Sea and Adriatic Sea) failed to reveal genetic differentiation (Passamonti et al., 1997). Maltagliati et al. (1998) performed allozyme analyses with fifteen enzyme systems in *N. norvegicus*, examining one Atlantic and eight Mediterranean samples, with approximately 100 individuals from each site. While genetic variability was detected, there was no evident population structure. Stamatis et al. (2006) used ten allozymes systems to investigate samples from the North Sea and the Aegean Sea, finding no significant genetic differentiation among 366 examined individuals. Streiff et al. (2001) did not recover evidence for population structure among forty individuals from two Portuguese locations for five microsatellite loci. Stamatis et al. (2004) performed a restriction fragment length polymorphism analysis on mitochondrial COI DNA segments in 370 individuals, and reported significant but low levels of genetic differentiation, but no structure between the Mediterranean Sea and Atlantic Ocean. Recent population expansion after the Last Glacial Maximum (LGM) was proposed as an explanation. Similarly, no population genetic structure was found using twelve microsatellite loci on 549 individuals from a small geographic range around Iceland (Pampoulie et al., 2011).

Previous studies have yet to recover population differentiation either across the geographic range of *N. norvegicus*, or at finer scales. The mtDNA D-loop has proven hypervariable in other crustacean species with high levels of polymorphism that can be used to discriminate amongst populations (Mcmillen-Jackson and Bert, 2003; 2004). The current study explores the efficacy of this region to determine the presence of population structure across a subsample of the species’ distribution.

## Materials and Methods

### Sampling

Samples were collected from commercial fishing or research vessels from across the geographic distribution of *N. norvegicus*, including Iceland, Northern Norway, Skagerrak, North Sea, Irish Sea, Porcupine bank, Bay of Biscay, Gulf of Cadiz and Ancona in the Adriatic Sea (Figure 1). Sex and length were recorded, and first and second pereiopods were removed from each individual before being stored in 807 EtOH. Whole samples that were collected were stored at -20 °C before tail tissue was removed and stored in 807 EtOH. Both males and females (~2:1) with carapace lengths encompassing an equal number (n=15) of individuals of two length groups (6-35mm & 35-70mm) were selected to minimise the risk of only including a single cohort that could cause family effects and skew the genetic data (Haynes et al., 2016).

### DNA Analysis

Total genomic DNA was extracted using a modified chloroform/isoamyl alcohol protocol (Petit et al., 1999). Primers (NN3DF 5′-ACA GCG TTA AGA YAC CAT AG-3′ and NnDR 5′-GCT CTC ATA AAC GGG GTA TGA-3′) were designed initially using Primer-3 as implemented in Geneious^®^ 7 (https://www.geneious.com, Kearse et al., 2012) and the D-loop *N. norvegicus* mitochondrial genome (GenBank Accession: LN681403.1). The resulting amplicons were relatively larger (~880bp) than had been designed for (~600bp) and were sequenced to discover a ~280bp fragment of the D-loop area missing from within the GenBank data (Appendix 1). Subsequently, new primers JG2 F 5′-CTA CAG ATT TCG TCT ATC AAC-3′ and NnD R 5′-GCT CTC ATA AAC GGG GTA TGA-3′ were designed on these returning sequences to incorporate the newly discovered ~280bp for a ~680bp amplicon. Primer sequences specificity was confirmed using BLAST (Basic Logical Alignment Search Tool; Zhang et al., 2000). Optimal annealing temperature was determined using a gradient PCR.

PCR amplifications were performed in a Biometra T3000 thermocycler (Biolabo, SA) with lid temperature of 95°C, using a thermal cycling profile of initial heating of 95°C for 2min, followed by 30 cycles of 95°C for 30s, 61.2°C for 30sec, and 72°C for 1min, followed by a final extension step of 72°C for 2min. Completed reactions were held at 4°C. PCR products were visualised on 1.57 agarose gels to verify amplifications, and purified using ExoSap-IT (Affymetrix Ltd., Santa Clara, CA) prior to Macrogen sequencing.

### Data and Statistical analyses

Forward and reverse sequences were aligned and edited in Geneious^®^ 7.0, using the K80 substitution model (Kimura, 1980), as determined in JMODELTEST, version 2.1.4. (Guindon and Gascuel, 2003; Darriba et al., 2012). Spatial Analysis of Molecular Variance, SAMOVA 2.0 (Dupanloup et al., 2002) was used to define groups of populations which are geographically homogenous and maximally differentiated from each other. Analysis of Molecular Variance (AMOVA) (Excoffier et al., 1992) was performed in ARLEQUIN V3.5 (Excoffier and Lischer, 2010) to generate F-statistics, Φ_ST_, θ, and (τ), the K80 substitution model and 10100 replicates. Fu’s F_s_ tests whether mutations are selectively neutral. Theta (θ) is defined as 2Nµ for haploid mitochondrial DNA, where N is the effective population size and µ is mutation rate per sequence per generation (Fu, 1997). Tau (τ) can measure relative time since a population expansion using T = τ/2u, where u is per-nucleotide rate of mutation multiplied by the number of nucleotides in the sequence (Gaggiotti and Excoffier, 2000). Harpending’s raggedness index (Harpending, 1994) and mismatch distributions *(SDD*) were both used to test whether the data deviated significantly from a population expansion model. DNASP version 6.10.01 (Rozas et al., 2017) was used to calculate haplotype (*h*) and nucleotide diversity (π) and to estimate and the nearest neighbour statistic S_nn_ (Hudson, 2000) with 10000 permutations. This statistic uses a symmetric island model on haplotype data to measure sequential ‘neighbours’ from the same geographical space. In all cases involving multiple comparisons, significance levels were adjusted for multiple tests using the sequential Bonferroni correction technique (Rice, 1989). A map of *N. norvegicus* haplotypes was constructed using POPART (Leigh and Bryant, 2015). An Unweighted Pair Group Method with Arithmetic Mean (UPGMA) dendrogram of Φ_ST_ pairwise distance values was created in PAST (Hammer et al., 2001). All software was used with default setting unless specified otherwise.

## Results

### DNA Analysis

Sequence alignments were trimmed for maximum length and quality using individual sequence chromatograms. Only regions of the D-loop for which both forward and reverse strands yielded unambiguous sequences were included for a fragment of 375bp. Of the 270 sequenced samples, 13 were excluded due to poor quality sequence reads. A total of 15 haplotypes were resolved (GenBank Accession: xxxxxx) with nucleotide diversity (π) ranging from 0.008 in the Gulf of Cadiz to 0.019 in the Bay of Biscay, and haplotype diversity ranged from 0.508 in Breiðamerkurdjúp to 0.859 in the Bay of Biscay (Table 1). Frequency and location of the haplotypes were displayed on a haplotype map (Figure 2A).

**Figure 2 A and 2 B.**
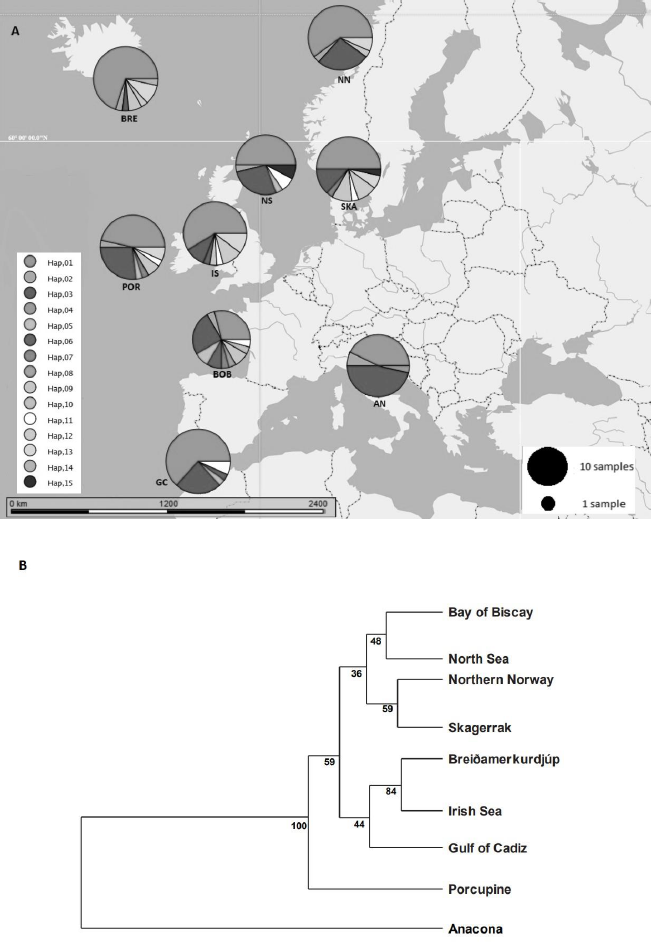
**2A:** Map of the geographic distribution and frequency of haplotype groups in nine sample sites of *N. norvegicus*. AN: Anacona, BOB: Bay of Biscay, BRE: Breiðamerkurdjúp, GC: Gulf of Cadiz, IS: Irish Sea (West), NN: Northern Norway, NS: North Sea, POR: Porcupine, SKA: Skagerrak haplotypes. **2B:** UPGMA dendrogram based on Φ_ST_ pairwise distance values calculated from frequency data of 15 haplotypes.

**Table 1.** Mitochondrial sequence variability in the D-loop for *N. norvegicus* from the nine sample sites; number of individuals (*n*), number of haplotypes (*nh*), haplotype diversity (*h*), nucleotide diversity (π), tau (τ), theta for times 0 and 1 (θ_0_ and θ_1_), Fu’s F_S,_ Harpending’s Raggedness index (*Hri*), and sum of squared differences from mismatch analyses (SSD)

| Collection | *n* | *nh* | *h* | $\pi$ | $\tau$ | $\theta_0$ | $\theta_1$ | Fu's $F_s$ | *Hri* | SSD |
| --- | --- | --- | --- | --- | --- | --- | --- | --- | --- | --- |
| Anaconda | 28 | 4 | 0.616 | 0.009 | 2.5 | 0 | 11.055 | **-26.691\*** | 0.040 | 0.006 |
| Bay of Biscay | 24 | 10 | 0.859 | 0.019 | 2.2 | 0 | 6829.96 | **-26.832\*** | **0.062** | 0.006 |
| Breiðamerkurdjúp | 30 | 7 | 0.508 | 0.010 | 2.8 | 0 | 6834.96 | **-26.748\*** | **0.055** | **0.010** |
| Gulf of Cadiz | 30 | 5 | 0.556 | 0.008 | 1.6 | 0 | 3429.40 | **-27.522\*** | **0.079** | 0.006 |
| Irish Sea | 29 | 7 | 0.643 | 0.018 | 1.6 | 0 | 3424.39 | **-27.717\*** | **0.138** | **0.022** |
| Northern Norway | 30 | 6 | 0.582 | 0.009 | 1.7 | 0 | 6844.37 | **-27.165\*** | **0.090** | **0.010** |
| North Sea | 26 | 5 | 0.689 | 0.012 | 2 | 0 | 6829.96 | **-26.775\*** | **0.091** | **0.011** |
| Porcupine | 30 | 8 | 0.722 | 0.013 | 2.5 | 0 | 6834.96 | **-26.647\*** | **0.046** | 0.003 |
| Skagerrak | 30 | 8 | 0.729 | 0.015 | 3 | 0 | 3414.98 | **-26.344\*** | **0.050** | **0.007** |

Values in bold are significant at $P < 0.05$ , \* $P < 0.001$

### Data and Statistical analyses

#### Population Differentiation

The SAMOVA analysis indicated that the best-supported F_CT_ value (F_CT_ = 0.109) was achieved when the samples were clustered into two groups. The first group contained only individuals derived from the Mediterranean (Anacona) while the second group comprised of individuals from all other sampled areas (Table 2).

**Table 2.** SAMOVA results table for *N. norvegicus.* K corresponds to the number of populations. Optimal F_CT_ and groupings (K=2) are highlighted in bold.

| Groups (K) | Structure Recovered | $F_{CT}$ |
| --- | --- | --- |
| **2** | **[AN][BOB, BRE, GC, IS, NN, NS, POR, SKA]** | **0.1092** |
| 3 | [AN][BOB][BRE, GC, IS, NN, NS, POR, SKA] | 0.0799 |
| 4 | [AN][POR][BRE, GC, IS, NN, SKA][BOB][NS] | 0.0717 |
| 5 | [POR][BRE, IS][BOB][AN][GC, NN, NS, SKA] | 0.0661 |
| 6 | [BRE, IS][AN][GC, NN, NS][SKA][POR][BOB] | 0.0618 |
| 7 | [AN][BRE, IS][POR][NN][GC, NS][SKA][BOB] | 0.0620 |
| 8 | [SKA][AN][NS][GC][BOB][NN][BRE, IS][POR] | 0.0637 |

An AMOVA analysis, using the SAMOVA structure and the K80 distance model (Kimura, 1980) revealed significant heterogeneity among the nine samples (Φ_ST_ = 0.107, P<0.001). Within-sample variation accounted for 88.867 of the variance (F_CT_=0.100, P<0.001) (Table 3). Pairwise ΦST values revealed population structure between the eastern Mediterranean (Anacona) sample and each of the eight other samples from the North Atlantic (Table 4). Significant pairwise Φ_ST_ values ranged from 0.057 (Anacona/Porcupine) to 0.152 (Anacona/Breiðamerkurdjúp) (Table 4). The nearest-neighbour statistic (S_nn_) indicated a significant association between sequence similarity and geographical location (S_nn_=0.1250, P=0.032). A UPGMA dendrogram was constructed using sample pairwise Φ_ST_ values to visualise genetic distances (Figure 2B).

**Table 3.** AMOVA table of temporal and spatial genetic variation of *N. norvegicus* from sites sampled.

| **AMOVA** |  |  |  |  |
| --- | --- | --- | --- | --- |
| Source of variation | % of variance | Fixation indexes | F-statistics | *P* |
| Among groups | 10.00 | 0.126 | FSC | <0.001 |
| Among samples within groups | 1.14 | 0.111 | FST | 0.785 |
| Within populations | 88.86 | 0.100 | FCT | 0.000 |

**Table 4.** Pairwise Φ_ST_ estimates (below diagonal) for D-loop mtDNA data among *N. norvegicus* samples. P-values (upper diagonal) in bold were significant after Sequential Bonferroni correction (initial α=0.05/8=0.00625).

| Population | Anaconda | Bay of Biscay | Breiðamer-kurdjúp | Gulf of Cadiz | Irish Sea | Northern Norway | North Sea | Porcupine | Skagerrak |
| --- | --- | --- | --- | --- | --- | --- | --- | --- | --- |
| Anaconda |  | **0.019** | **<0.001** | **0.021** | **0.024** | **0.031** | **0.046** | **0.026** | **0.016** |
| Bay of Biscay | 0.111 |  | 1.336 | 1.134 | 0.270 | 0.870 | 1.062 | 0.273 | 0.717 |
| Breiðamer-kurdjúp | 0.152 | 0.046 |  | 0.572 | 0.615 | 0.627 | 0.200 | 0.060 | 0.670 |
| Gulf of Cadiz | 0.144 | 0.002 | 0.019 |  | 1.196 | 1.434 | 1.356 | 0.140 | 0.704 |
| Irish Sea | 0.143 | 0.043 | -0.009 | 0.005 |  | 0.403 | 0.333 | 0.240 | 0.604 |
| Northern Norway | 0.072 | 0.018 | 0.011 | -0.006 | -0.001 |  | 1.248 | 1.554 | 0.748 |
| North Sea | 0.087 | -0.007 | 0.043 | -0.015 | 0.027 | -0.010 |  | 0.358 | 0.625 |
| Porcupine | 0.057 | 0.049 | 0.061 | 0.051 | 0.037 | -0.010 | 0.015 |  | 0.129 |
| Skagerrak | 0.098 | -0.014 | 0.002 | -0.010 | -0.008 | -0.010 | -0.009 | 0.020 |  |

#### Population Expansion

Demographic analyses in ARLEQUIN showed pronounced differences between θ_0_ and θ_1_ suggesting rapid population expansion in all samples, with less pronounced differences in the Anacona sample (Table 1). All Fu’s F_s_ values were negative and deviated significantly from zero. Mismatch distributions differed significantly from the distributions expected under population expansion in four of the nine sample sites (Table 1). Harpending’s raggedness index ranged from 0.040 to 0.138 from the Anacona to the Irish Sea respectively and was significant for all except the Anacona sample (Table 1).

Using the population expansion formula T=τ/2u, with u=μk, where μ=per nucleotide substitution rate and k= sequence length population expansion times were estimated between 10,500 to 19,000 ybp. Due to the uncertainty around the point estimate of τ in the Arlequin analyses, 1000 bootstrap replicates were performed drawing random values for tau from between 2.5 and 97.5 percentiles returned by Arlequin. Population expansion times were estimated using μ=197/My from the penaeid prawn and pink shrimp D-loop mutation rate (Mcmillen-Jackson and Bert, 2003; 2004). From the 1000 bootstrap iterations the mean estimate and upper and lower two standard deviation confidence intervals were calculated (Table 5).

**Table 5.**
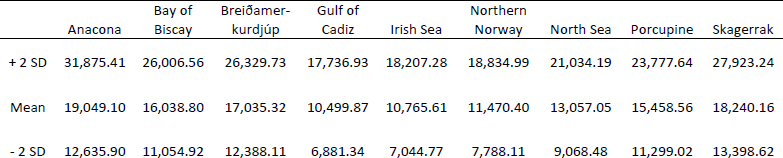
Estimated expansion times for *N. norvegicus* with upper and lower two standard deviation confidence intervals.

|  | Anaconda | Bay of Biscay | Breiðamer-kurdjúp | Gulf of Cadiz | Irish Sea | Northern Norway | North Sea | Porcupine | Skagerrak |
| --- | --- | --- | --- | --- | --- | --- | --- | --- | --- |
| + 2 SD | 31,875.41 | 26,006.56 | 26,329.73 | 17,736.93 | 18,207.28 | 18,834.99 | 21,034.19 | 23,777.64 | 27,923.24 |
| Mean | 19,049.10 | 16,038.80 | 17,035.32 | 10,499.87 | 10,765.61 | 11,470.40 | 13,057.05 | 15,458.56 | 18,240.16 |
| - 2 SD | 12,635.90 | 11,054.92 | 12,388.11 | 6,881.34 | 7,044.77 | 7,788.11 | 9,068.48 | 11,299.02 | 13,398.62 |

## Discussion

This study recovered a previously-undocumented 280bp segment of the *N. norvegicus* mitochondrial genome (GenBank Accession: xxxxx). In addition, genetic structure between North Atlantic and Mediterranean Sea *N. norvegicus* samples was detected in the mtDNA D-loop region. While an Atlantic-Mediterranean divide has been recorded for many highly mobile species (Bargelloni et al., 2003; Carlsson et al., 2004; Farrell et al., 2016), it has not been previously reported for *N. norvegicus*.

Eleven haplotypes are shared among multiple *N. Norvegicus* samples, with four haplotypes unique to a single sample: two unique haplotypes are found in the Bay of Biscay, and one each are found in Breiðamerkurdjúp and Anacona. SAMOVA revealed distinct population genetic differences between the Atlantic samples and the eastern Mediterranean sample, and the nearest neighbour statistic (S_nn_) suggests a significant association between D-loop sequence similarity and geographic location. Average Φ_ST_ estimates are at least twice as large between the Mediterranean sample and each Atlantic sample than the estimates among all of the Atlantic samples. This suggests that the eastern Mediterranean sample is genetically differentiated from the Atlantic samples. A UPGMA cluster analysis on the Φ_ST_ distance matrix demonstrated that the largest genetic differentiation exists between the Mediterranean and all other samples.

Negative Fu F_s_ values suggest recent demographic expansion (Fu, 1997), and the large difference between θ_0_ and θ_1_ for all Atlantic sites suggests rapid population expansion. In contrast, the difference in theta values for the Mediterranean sample are two orders of magnitude smaller. When considered with the non-significant Raggedness index, this suggest a less pronounced population expansion in the Mediterranean.

Estimates of time since expansion ranged from 10,500 to 19,000 ybp. Large confidence intervals around all of the point estimates for the expansion time overlap, indicating that expansion likely occurred within the same time-frame for all sampled locations. These time estimates are in agreement with those for the LGM in Europe (16,000 to 31,000 ybp, Ashton et al., 2010), and likely represent population expansion into newly available habitat as the ice retreated. Observed haplotype diversity was highest in Bay of Biscay, suggesting this region represents a potential glacial refugium for the Atlantic distribution of the species. The area north of the Bay of Biscay has previously been hypothesized as a refugium for other marine species (e.g., the common mussel *Mytilus edulis*, (Linnaeus, 1758), Śmietanka et al., 2014), and these results also support species distribution models for several other marine invertebrates, including the common starfish, *Asterias rubens* (Johnston, 1836), amphipod crustacean *Gammarus duebeni* (Liljeborg, 1852), flat periwinkle *Littorina obtusata* (Linnaeus, 1758), dogwhelk *Nucella lapillus* (Linnaeus, 1758) and barnacle *Semibalanus balanoid* (Linnaeus, 1767) around the LGM (Waltari and Hickerson, 2013).

This study is the first to reveal a significant genetic differentiation between Atlantic and Mediterranean samples of *N. norvegicus*. These results support a post-glacial expansion, with Atlantic *N. norvegicus* continuing to expand rapidly. In terms of commercial fisheries management, these results do not support current management practices, as no significant genetic differentiation was found among Atlantic samples that cross several functional units. However, these results are important for management within the eastern Mediterranean, as populations experiencing isolation can be more vulnerable to commercial over-exploitation and more difficult to recover in the event of population collapse.

## Acknowledgements

We would like to thank the scientists and researchers from the Irish Marine Institute, Ifremer, Marine Research Institute Iceland, Spanish Institute of Oceanography, ISMAR Institute of Marine Sciences, Havforskningsinstituttet *Norwegian Institute of Marine Research*, Marine Scotland and AFBI for the collection and donation of samples involved in this study. We would also like to thank E. Farrell for his assistance and comments on the manuscript.

## Appendices

### Appendix 1

*Nephrops norvegicus* D-loop sequence (GenBank Accession: xxxxx) with ∼280bp fragment (shown in bold) missing from within the GenBank data.

ATATACACAGATCAGTAAAAATATATTTTTAAGGCTAATCTAAAAAGTAAACTTATATAATTTCATT GAAATTCATTACARTCTGAAAGTCAATGATTTAATTTTATAAATCGACTAAATAAGATCTATAAATA AAA**TCTTACCCCTTCAAAAGGTCACTTTCTCCTGAGGGGAGCTCCCTTTTCCCAACGGGGTAAGATT TCTATTGGGAGAGCAGGATTATAATTATAGAGAGTTGGGTATAAGGCTTCATTGTTTACACATATAT ACTATTAAATTAATTATATACATTTATATGTATATATATATATATATATATACTATTTAAATAATAT TTTCTTAACTTTWTATTTTGTTAACATWTAAATTATTAATAATGTTTTATAAATTTTATATATTAAA ATAAAATACAGTAAAA**AAGGTTTTTAGATAAATTTCTACGAATATTATACTATTATACACAATGGAA TTCCACCAATTCTTTAAAGATCAAAACTTTTCGTGCCGTTTACACTAGTATACAAAAGAGAAGCTAA TTCTAAGCTAATGG

